# DARE: Division Axis and Region Estimation from 2D and 3D Time-Lapse Images

**DOI:** 10.1101/2024.02.05.578987

**Authors:** Romain Karpinski, Alice Gros, Marc Karnat, Qazi Saaheelur Rahaman, Jules Vanaret, Mehdi Saadaoui, Sham Tlili, Jean-François Rupprecht

## Abstract

We propose a two-stage supervised framework for characterizing cell divisions in 2D and 3D time-lapse microscopy. First, we recast division detection as a semantic segmentation task on image sequences. Second, a local regression model estimates the orientation and distance between daughter cells for each event. We validate this framework using image sequences of avian neuroepithelium and mouse gastruloids. Our results demonstrate that high performance is achieved with efficient architectures, namely a U-Net for segmentation and a CNN for regression, that are optimized through systematic hyperparameter exploration. We find that integrating temporal context via multiple consecutive frames significantly boosts segmentation accuracy. We achieve F1 scores exceeding 94% (2D+t) and 90% (3D+t), with orientation accuracy approaching the uncertainty limit of manual annotation. We provide the full codebase and training workflow, specifically designed for datasets where traditional tracking is challenging.

## I. INTRODUCTION

Identifying cell division events is essential for understanding tissue dynamics. While static markers like Ki-67 or EdU provide a census of proliferating cells, they fail to capture the spatiotemporal correlations necessary to understand how divisions interact over time [1, 2]. These dynamics are central to tissue mechanics, where division serves a dual purpose: generating mechanical stress through mitotic rounding [3] and relaxing it through topological remodeling [4, 5]. Mapping these events relative to tissue wounding, crowding, or signaling gradients is therefore critical for deciphering the feedback loops that maintain homeostasis [6, 7].

However, robustly extracting these events from live-imaging data remains a bottleneck. Traditional lineage reconstruction depends on high-fidelity segmentation across all time points [8–10], a requirement that is rarely met in dense, scattering 3D live animal tissues, which are often plagued by optical aberrations and stringent photon-budget constraints, both of which severely degrade segmentation and tracking performance. Marker choice only partially alleviates these issues: nuclear labels can simplify mitosis detection but provide limited information on cell boundaries and topology, while membrane or contour signals better capture tissue geometry and mechanics, they are often harder to segment robustly, especially in 3D. In these challenging conditions—characterized by anisotropic resolution and rapid neighbor exchanges—global tracking often becomes unreliable [11].

Detecting divisions as discrete point events offers a significant advantage, because they are associated with distinct visual signatures, such as sharp chromatin condensation followed by rapid changes in cell geometry. Capitalizing on such visual signatures, several recent computational frameworks have shifted toward the direct detection of division events within image sequences. McDole et al. [2] were the first to use a U-Net-like segmentation network with multi-frame input (*N* = 7 frames) for the detection of division centers. However, their framework was not designed to infer the orientation of daughter cells, which prevented the extraction of division angles. More recently, Turley et al. [12] combined two complementary temporal U-Nets: one dedicated to localizing division events in 2D sequences and a second to inferring orientation via a segmentation mask. However, this latter step requires additional image post-processing to determine the final angle.

In this work, we evaluate a two-stage location/orientation strategy conceptually similar to [12], but with a key architectural change: we replace the second temporal U-Net with a dedicated CNN that directly regresses the division angle and length, see Fig. 1. This approach eliminates the post-processing step required in Ref. [12] to extract the distance and orientation of the daughter cells from the U-Net prediction map. Elements of our methodology, first introduced in [13], had already been applied in [14]. Furthermore, we systematically investigate the impact of several hyperparameters, including the input frame count (*N*), on algorithmic performance. Finally, we extend our framework to challenging 3D image sequences, demonstrating that it achieves robust performance with only a few hundred human annotations—a significant reduction compared to the data requirements typically seen in 2D contexts.

**FIG. 1.**
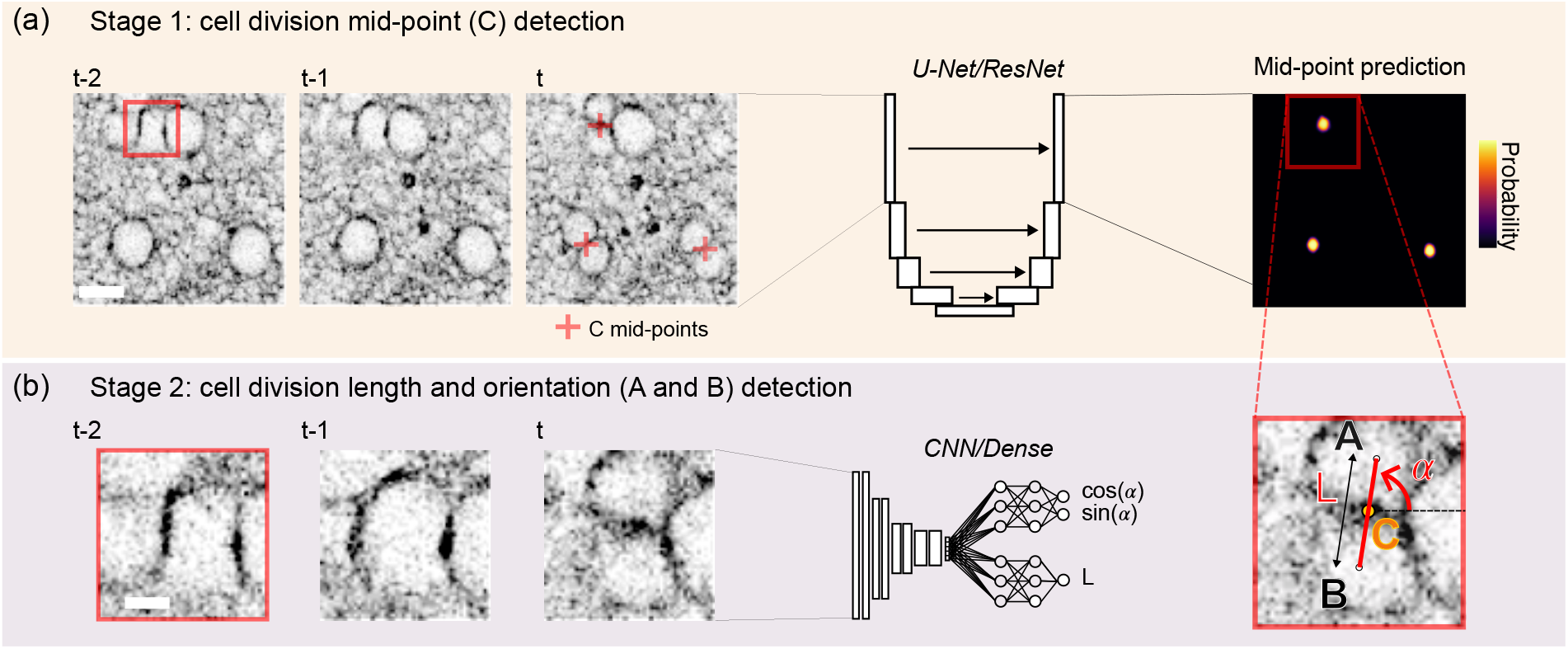
Description of our two-stage approach, used across all models (illustrated here in 2D on the avian neuroepithelium Dataset 1). (Top row) Stage 1: the input image sequence is processed by a U-Net to predict division centers. The image sequence is stacked along the channel dimension. Scale bar: 10 *µ*m. (Bottom row) Stage 2: for each predicted division center, a local patch is cropped and fed to a regression CNN to estimate the division length and orientation. The regressor also uses temporally stacked input images from multiple time steps. Scale bar: 3 *µ*m.

By extracting the timing and orientation of divisions independently of continuous tracking, our cell-event detection framework could be integrated within tissue-scale measures — such as the estimation of flow fields (through particle image velocimetry or optical flow) — to bridge the gap between single-cell behavior and collective tissue mechanics [15, 16].

## II. DATASETS

Here, we consider the following datasets:

- Datasets 1 and 2: avian neuroepithelium (neural tube) explants labeled with a membrane marker (SiR-actin). Neural cells first migrate along the *z*-stack direction (called the apicobasal axis) before dividing [3]. Using inverted confocal microscopy (Zeiss LSM 880/1.1 C-Apochromat objective), 3D time-lapse (image stacks) were acquired every 3 min. (Fig. 1). Within each *z*-stack, each pixel corresponds to 1.2 *µ*m, and dividing cells have a typical diameter of ≈ 10 *µ*m. In Dataset 1 (2D), each set consists of a time sequence acquired at a particular *z*-plane. In Dataset 2, each volume consists of sequential optical planes spaced by 1 *µ*m. Within the fixed observation planes used to generate Dataset 1, cells first appear as growing disks that, upon reaching a critical size, undergo fission: the cell membrane pinches along a diameter, eventually splitting into two.
- Dataset 3 (3D): mouse gastruloids, i.e. 3D self-organizing aggregates derived from embryonic stem cells that recapitulate key aspects of early gastrulation, including axial patterning and germ-layer specification [17, 18]. Using two-photon microscopy (Zeiss 510 NLO inverted LSM, femtosecond laser Mai Tai DeepSee HP, 40*×* /1.2 C-Apochromat objective), 3D time-lapse (image stacks) were acquired every 4 min. Each volume consists of sequential optical planes spaced by 1 *µ*m. Nuclei typically occupy a volume of ∼ 5*×*5*×*5 *µ*m^3^. The gastruloids were formed from genetically modified cell lines expressing the fluorescent nuclear marker H2B–GFP, as in [19]. The H2B–GFP signal was excited at 920 nm and detected with a non-descanned GaAsP detector using a 560 LP filter. Imaging depth in *z* ranged from 50 to 150 *µ*m, depending on acquisition settings and sample size. Gastruloids were imaged at multiple developmental stages. At 72 h after formation (**Movie** 4), they remain largely round and exhibit highly dynamic cell migration and frequent neighbor exchanges. At 96 h after formation (**Movie** 5–**Movie** 6), they are elongated, display gradients of gene expression along the elongation axis, and show slightly slower cell dynamics together with stronger spatial heterogeneity in cell density.

### Annotations

For each division, the annotations consist of two disks (resp. spheres) of radius 7 pixels (resp. voxels) in 2D (resp. 3D) centered on the two daughter cells (resp. nuclei) in the first frame in which they become visible; in the nuclei/gastruloid context, the annotated frame corresponds to the first frame after chromosome separation (telophase). Annotations were performed using napari [20].

Divisions were particularly challenging to annotate when oriented perpendicular to the imaging plane, i.e. along the least-resolved axis (Fig. 2b). To facilitate 3D annotation, we employed depth color coding of z-planes through the napari plugin `napari-zplane-depth-colorizer` [21]; we applied it in its default mode for **Movie** 4–**Movie** 6. Independent cross-checking between experts was used to minimize errors in the ground-truth dataset.

**FIG. 2.**
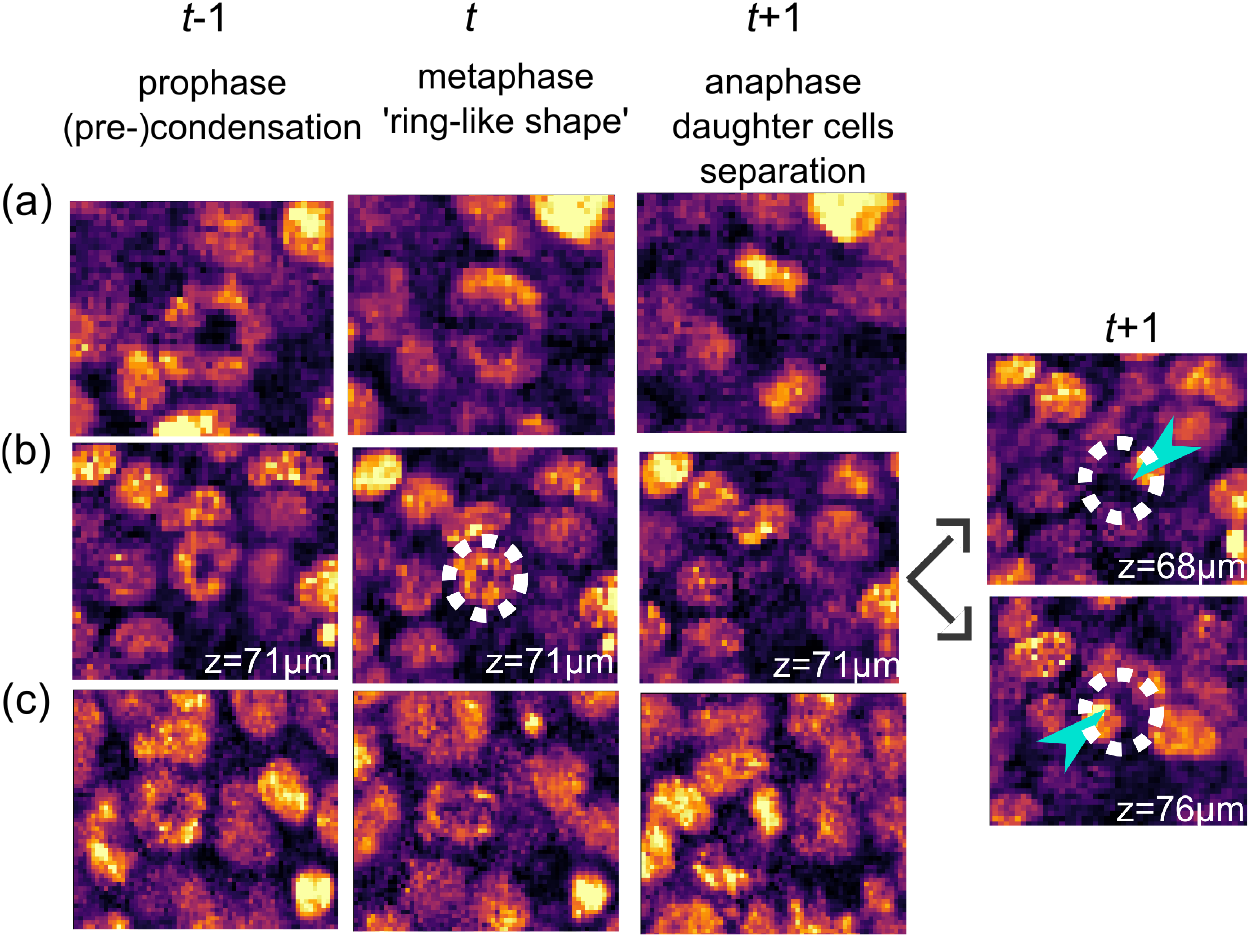
Dataset 3: gastruloids, nuclei signal (histone GFP). Live division detection is challenging in 3D, as illustrated by three cases of increasing complexity, from (a) to (c), shown across three consecutive frames (columns *t* − 1, *t, t* + 1; see Methods section, DARE3d network input). (a) The easiest case to annotate is when both the metaphase ring-like structure and the two daughter cells lie within a single *z*-slice. (b) A more complex case occurs when the ring structure lies clearly within a single *z*-slice, but the two daughter cells do not. Blue arrows indicate the two daughter cells, and the dotted circle shows the position of the mother cell at separation time *t* + 1. (c) Complex cases include situations in which daughter cells have reduced brightness and contrast compared to the surrounding cells.

## III. METHODS

Here we introduce the Division Axis and Region Estimation (DARE) method. It is implemented as DARE2d (a TensorFlow-based framework) and DARE3d (a PyTorch-based framework).

### A. Methods: 2D

#### 1. Stage 1: midpoint segmentation by an N-frame U-Net

##### Cell division midpoint segmentation radius r

We approach cell-division detection as a semantic segmentation task. We initially considered representing the two daughter cells, either by individual centers or by an ellipse spanning them; however, pairing daughter cells becomes ambiguous in dense division clusters. We therefore represent each division by its midpoint (point *C* in Fig. 1a), i.e. the center of the cleavage furrow. This center is encoded as a circular region of radius *r*.

##### Network architecture

We use a U-Net architecture [22] with a ResNet-18 encoder [23], followed by five successive upsampling and convolution blocks in the decoder. The final activation function is a sigmoid.

##### Input processing

We use as input image sequences of size 256 *×* 256 pixels, with *N* ∈ {1, 2, 3} consecutive frames stacked along the channel dimension. Images are pre-processed using histogram equalization. Since the task is binary with a strong class imbalance (many background pixels), we assign a weight *W* to the division-midpoint class in a weighted binary cross-entropy (WCE) loss:

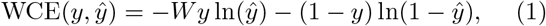

where *y* denotes the ground truth (*y* = 1 on division-midpoint pixels and 0 otherwise), and *ŷ* denotes the predicted probability.

##### Data augmentation and training

We use a standard combination of random brightness changes, flips, Gaussian noise, rotations, elastic deformations, and random scaling using the Albumentations library [24], as well as intensity-histogram deformations [25]. We train the network for 75 epochs, with 250 iterations per epoch and a batch size of 32, using the Adam optimizer with a learning rate of 10^−4^.

##### Post-processing

We apply a threshold to the output probability map (*P >* 0.5). The centroids of the resulting connected components define the predicted midpoint positions. In some instances, the same division is detected in two consecutive frames; such redundancies are filtered by comparing centroid distances across successive frames.

##### Evaluation metrics

Because daughter-cell separation can span multiple frames, a *±*1 frame tolerance is used. To count true positives, we apply a two-step matching scheme: at each time *t*, (1) we match predicted centers to the ground-truth set at time *t* within a spatial tolerance of 10 pixels (12 *µ*m), and (2) unmatched predicted centers are then matched to the ground-truth sets at times *t* − 1 and *t* + 1. Predicted centers that remain unmatched are counted as false positives (FP), and ground-truth centers that remain unmatched are counted as false negatives (FN). We then compute the F1 score

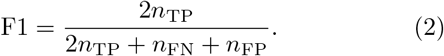

with *n*_TP_, *n*_FN_, and *n*_FP_ the numbers of true positives (TP), false negatives (FN), and false positives (FP), respectively.

#### 2. Stage 2: orientation and length by a CNN regressor

##### Input

We use as input a crop of size 64 *×* 64 pixels centered on the detected division midpoint. This size is chosen to encompass the two daughter cells, as the inter-daughter distance typically lies in the 20–40-pixel range.

##### Architecture

We use a CNN composed of four convolutional blocks. Each block comprises a convolution layer followed by a ReLU activation and a max-pooling layer. The first block uses 128 filters, and this number is doubled in each subsequent block. After the fourth block, we use two regression heads on the flattened features: one scalar *l* for the length, and one vector (cos(2*α*), sin(2*α*)) for the orientation. This representation is preferred over the scalar *α* to avoid the discontinuity between 0 and *π*, which correspond to the same orientation but are far apart under a standard MSE loss [26]. We use data augmentation and training protocols similar to those described for the U-Net.

#### 3. Post-processing: DARE2d ensemble model

In the 2D setting, the dataset comprises a large number of annotated divisions distributed across eight independent experimental movies (Dataset 1; Table I). This enables an experiment-wise *k*-fold cross-validation strategy with *k* = 8: in each fold, a model is trained on seven movies and evaluated on the held-out movie, yielding *k* = 8 independently trained models. This enables us to form an ensemble consensus from the predictions of these *k* = 8 models (bagging-style aggregation [27]), which suppresses outliers, removes double counting, and provides uncertainty estimates. Candidate divisions are clustered in space (and within a *±*1-frame temporal window) using density-based clustering (DBSCAN with proximity radius *ε* = 10 pixels) on the predicted midpoints. If a cluster contains at least 6 points (out of 8), it is considered a single detected division, with location and orientation given by the cluster average and the confidence interval given by the cluster dispersion, reported as the *±σ* (standard deviation) in **Movie** 2, **Movie** 3. This procedure defines the DARE2d ensemble model.

**TABLE I.** Description of Datasets 1–3: marker type (membrane/nuclei), age of sample (*A*, 72h or 96h), image/stack size in pixels (px), number of frames (Δ*T*), number of annotated divisions (*N*_div_), and mean division length in pixels (*l*) for each set. For Dataset 3, set 1 corresponds to **Movie** 4 and set 2 to **Movie** 5. The entries highlighted in blue denote the test subsets of model 8 for Dataset 1, and of the models provided for Datasets 2 and 3.

| Dataset | Signal | A | Set | Size (px) | $\Delta T$ | $N_{\text{div}}$ | $l$ (px) |
| --- | --- | --- | --- | --- | --- | --- | --- |
| 1 | memb. | 72h | 1 | (768,768) | 33 | 585 | $23.5 \pm 0.1$ |
| | | | 2 | (768,768) | 100 | 226 | $37.4 \pm 0.3$ |
| | | | 3 | (1024,1024) | 84 | 706 | $36.8 \pm 0.2$ |
| | | | 4 | (768,768) | 100 | 623 | $23.2 \pm 0.1$ |
| | | | 5 | (1024,1024) | 186 | 1272 | $33.4 \pm 0.1$ |
| | | | 6 | (1024,1024) | 186 | 942 | $33.6 \pm 0.1$ |
| | | | 7 | (1024,1024) | 56 | 445 | $35.9 \pm 0.2$ |
| | | | 8 | (1024,1024) | 55 | 279 | $36.0 \pm 0.2$ |
| 2 | memb. | 72h | 1 | (1024, 1024, 10) | 26 | 226 | $29.6 \pm 0.3$ |
| | | | 2 | (1024, 1024, 10) | 21 | 145 | $32.5 \pm 0.3$ |
| | | | 3 | (1024, 1024, 10) | 11 | 104 | $31.3 \pm 0.3$ |
| 3 | nuclei | 72h | 1 | (362, 305, 180) | 10 | 226 | $15.3 \pm 0.2$ |
| | | 96h | 2 | (424, 411, 85) | 28 | 210 | $13.0 \pm 0.2$ |
| | | 96h | 3 | (405, 394, 152) | 28 | 316 | $12.8 \pm 0.1$ |

### B. Methods: 3D

#### 1. Stage 1: midpoint segmentation by a 3-frame 3D U-Net

##### Cell division midpoint segmentation radius r

As in the 2D case, we represent each division by the midpoint between the two daughter cells (point *C* in Fig. 1a). This center is encoded as a spherical region of radius *r* = 8 voxels. It approximately corresponds to the location of the mother cell in the frame preceding its separation into two daughter cells.

##### Data processing

We use as input image sequences of size 128 *×* 128 *×* 128 voxels, with *N* = 3 consecutive frames stacked along the channel dimension. Images are pre-processed using histogram equalization and the local contrast enhancement method developed in [19]. Volumes are resampled to an isotropic voxel size of 1 *µ*m in *x, y*, and *z* using first-order interpolation. During training, we use a balanced sampling strategy with 50% positive crops (containing at least one division) and 50% hard negatives (containing no divisions).

##### Network architecture and training

We use a 3D U-Net [28] with six stages comprising 32, 64, 128, 256, 512, and 1024 filters. Spatial resolution is reduced by a factor of two after each stage. Each stage uses three residual units and batch normalization. The network input consists of the three time points concatenated along the channel dimension. We train for 200 epochs using a onecycle learning-rate scheduler, with a batch size of 32. We use the Adam optimizer with an initial learning rate of *l*_*r*_ = 0.1. The loss is the Unified Dice–Focal loss, a linear combination of Dice and Focal losses [29].

##### Augmentation

We use random flips, random scaling, random rotations, random contrast changes (gamma transform), and random non-linear histogram shifts, using the MONAI augmentation toolbox [30].

#### 2. Stage 2: orientation and length by a CNN regressor

##### Input

We use as input a crop of size 32 *×* 32 *×* 32 voxels, centered on the detected division midpoint.

##### Output

The CNN regressor outputs both a rotation describing the division-axis orientation and a scalar length corresponding to the daughter-cell separation. For the rotation, we follow the approach initiated in Ref. [31]; in 3D, a rotation matrix can be written as

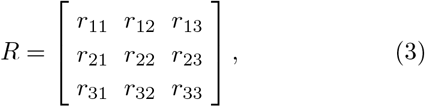

where the columns form an orthonormal basis. Regressing the nine entries of *R* directly does not in general yield an exactly orthonormal matrix. Instead, we regress a matrix that is orthogonalized within the loss, so that the corresponding rotation is well defined. Following [31], we use singular value decomposition (SVD) for orthogonalization.

##### Architecture

The regressor is composed of five 3D convolutional blocks (3D convolution + ReLU + 3D maxpooling), starting from 16 filters and doubling the number of filters at each stage.

#### 3. Post-processing

##### Thresholding

The probability map obtained after segmentation is first thresholded at a value *τ* selected on the validation set to maximize the F1 score. The resulting binary mask typically contains small spurious connected components. To remove those, for each connected component *O*, we compute (i) its mean probability

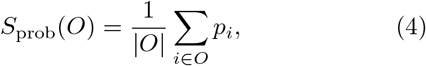

and (ii) a volume-consistency term

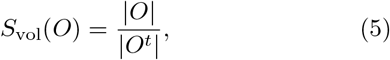

where |*O*| is the number of voxels in the component and |*O*^*t*^| is the expected number of voxels for the target object used during training (a sphere of radius *r* = 8 voxels). We then define a combined score

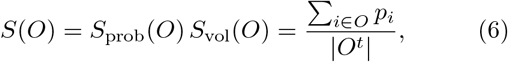

and retain a component as a valid detection only if *S*(*O*) *> τ*_*S*_, where *τ*_*S*_ is selected on the validation set.

##### Removing double-counting events

As in the 2D case, the same division can be detected in two consecutive frames. We therefore merge redundant detections by grouping components using a 4D distance criterion (10-voxel spatial tolerance and a *±*1 frame tolerance).

## IV. RESULTS

In both 2D and 3D, we use *N* = 3 input frames (see Discussion, Sec. V A).

### A. Results: 2D

We apply a *k* = 8-fold cross-validation procedure, where each fold corresponds to one independent experiment (Table I). In each fold, models are trained on seven experiments and evaluated on the held-out experiment, ensuring that no frames from the validation movie are seen during training.

Pooled across all validation folds, the method achieves an average F1 detection score of 0.955 *±* 0.015 (Fig. 3a– c), demonstrating both high accuracy and low variance across experimental conditions. The standard deviation of the orientation-angle error reached 3 *±* 0.5^°^ (see Discussion, Sec. V C).

**FIG. 3.**
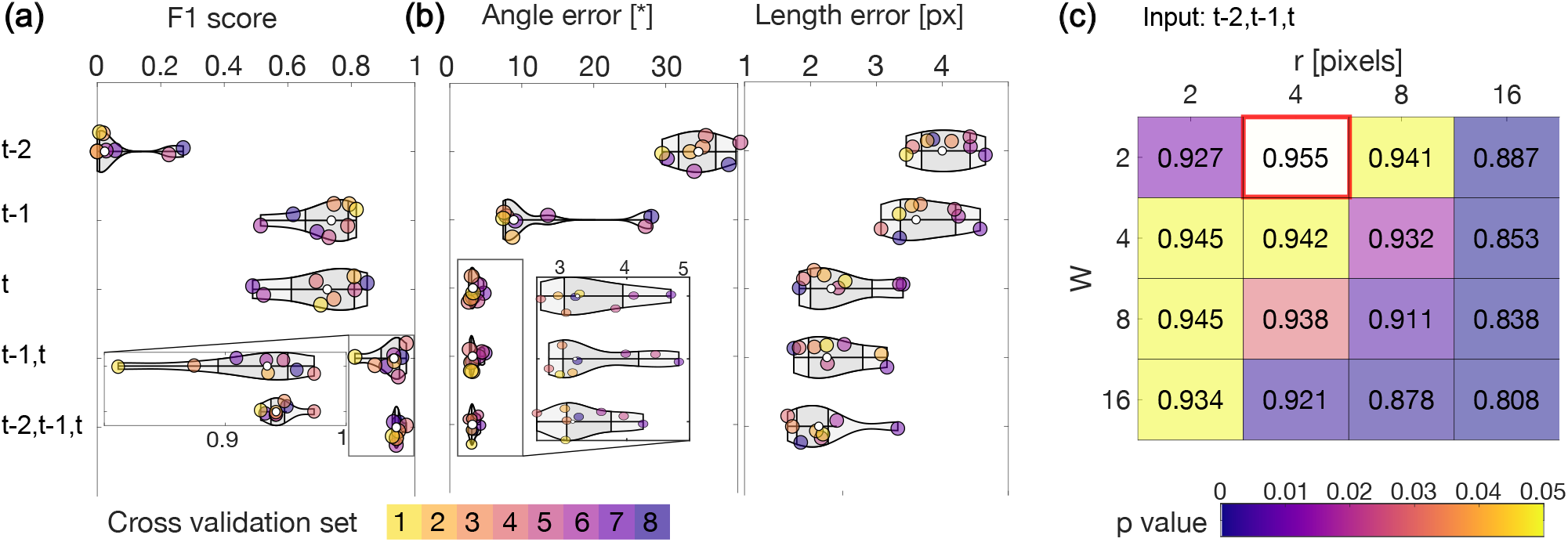
DARE2d membrane results and 8 models discussion: (a-b) Impact of the input time window on (a) the F1 score of the midpoint detection task, (b) the error on the orientation and length inference. The time *t* refers to the first frame at which membrane separation between the two daughter cells is apparent. Each colour refers to the set used for the validation. (c) Impact of the weight scaling *W*, defined in Eq. (1) and circle radius *r* on the mean F1 performance of the U-Net-based division location inference with (*t* − 2, *t* − 1, *t*) as input. The color code (shades of blue) refers to a *p*-value, estimated by a two-sample t-test, against the network’s best performance for *W* = 2 and *r* = 4px (red box). Yellow shading indicates that the p-value is above 5%, meaning that the difference from the optimal value is not statistically significant at this level.

### B. Results: 3D

#### 1. Results: 3D nuclei model

On our test (frames from set 1, Tab. I), the model achieves 124 TP, 1 FP, and 21 FN, corresponding to an F1 detection score of 91.8%. The standard deviation of the orientation-angle error is 28^°^ (see Discussion, Sec. V C).

#### 2. Results: 3D membrane model

On our test dataset, the model achieves 114 TP, 8 FP, and 8 FN, corresponding to an F1 detection score of 93.4% for division detection. In addition, the standard deviation of the orientation-angle error is 28^°^ (see Discussion, Sec. V C).

## V. DISCUSSION

### A. Discussion: 2D membrane model

#### Optimal (r, W)

To identify the optimal parameter set, we performed a grid search over the class weight *W* for division-midpoint pixels, see Eq. (1), and the target radius *r*, and selected the combination that maximizes the F1 score of our best *N* = 3-frame U-Net, see Fig. 3c. We find that the pair *W* = 2 and *r* = 4 px is optimal; increases in *W* can be compensated by decreases in *r*, see Fig. 3c.

#### Impact of the number of input frames

Using the two preceding frames (*t* − 2, *t* − 1, *t*) yields a significant improvement in the F1 detection score (0.945 *±* 0.015) for midpoint detection compared to the other time windows considered (e.g., 0.918*±*0.053 for (*t* − 1, *t*); Fig. 3a). The improvement is less pronounced for angle and length regression: compared to (*t* − 1, *t*), using (*t* − 2, *t* − 1, *t*) reduces the mean angle error and the mean length error by 0.4^°^ and 0.2 px, respectively (Fig. 3b).

#### Influence of ground-truth versus predicted centers

Regression performance is robust to errors in the detected midpoint location. Using ground-truth versus predicted midpoints changes the mean angle error by only 0.3^°^ and the mean length error by 0.7 px (model with *W* = 2 and *r* = 4 px).

#### Localisation error: FN/FP investigation

The fact that divisions can span more than two frames (Fig. 6a) explains why including the (*t* − 2) frame improves performance. In particular, using only *t* or (*t* − 1, *t*) can miss divisions that extend over several frames (Fig. 6a). To analyze the remaining errors of our best (*t* − 2, *t* − 1, *t*)-frame U-Net, we examined the 14 FN and 11 FP among the 268 ground-truth annotations in set 3, Table I, see Fig. 6. Importantly, the evaluation includes image-border regions (i.e., no margin was excluded). We find that 8 FN and 6 FP occur near the image boundary, where missing context and cropping likely explain the failures. Among the remaining FN, some correspond to clear divisions, but rare appearances—such as high membrane intensity within the cell bulk, see Fig. 6b —can reduce detection confidence. By contrast, almost all of the remaining FP correspond to ambiguous sequences that can confuse even experts. In particular, cells can rapidly migrate from the basal side (low *z*-planes) and enter the imaged apical plane (high *z*-planes); when this occurs near a growing cell, the resulting sequence resembles a division, producing FP (Fig. 6c). The F1-scores we obtain are also similar to the network with *N* = 10 frames as input from [12].

#### Orientation error

Angular accuracy is ultimately limited by the variability of manual annotations. A one-pixel uncertainty on each endpoint of a 25-pixel-long segment corresponds to an angular uncertainty of approximately (180*/π*) *×* (1*/*25) ≃ 2.3^°^. The mean angular error of DARE2d is not significantly larger than this value.

### B. Discussion: 3D

#### 1. Discussion: 3D nuclei model

##### Localisation error

Our DARE3d nuclei model successfully detects divisions across a range of orientations, including divisions perpendicular to the imaging plane and events in low signal-to-noise regions (Fig. 4a and **Movie** 6). We interpret several false positives (FPs) as dead cells (Fig. 7); indeed, dead cells share the following features with dividing nuclei in telophase: (i) increased brightness in the histone signal, (ii) rapid trajectories when dead cells are transported and cleared during tissue growth, and (iii) fragmentation into multiple debris, similarly to daughter-cell separation during telophase. The FP corresponds to a bright cell displaying an unusually fast trajectory and is potentially a dead cell. 10 false negative cases (FN) were divisions for which the probability map did not exceed the threshold; the remaining FNs were divisions that occurred very close to each other and were merged into a single detection during post-processing. The threshold value (accepting or rejecting a local maximum in the probability map as an event), as well as the minimum distance between two events, are two key parameters optimised to maximise the F1 score; both are accessible and can be tuned in the code for further applications.

**FIG. 4.**
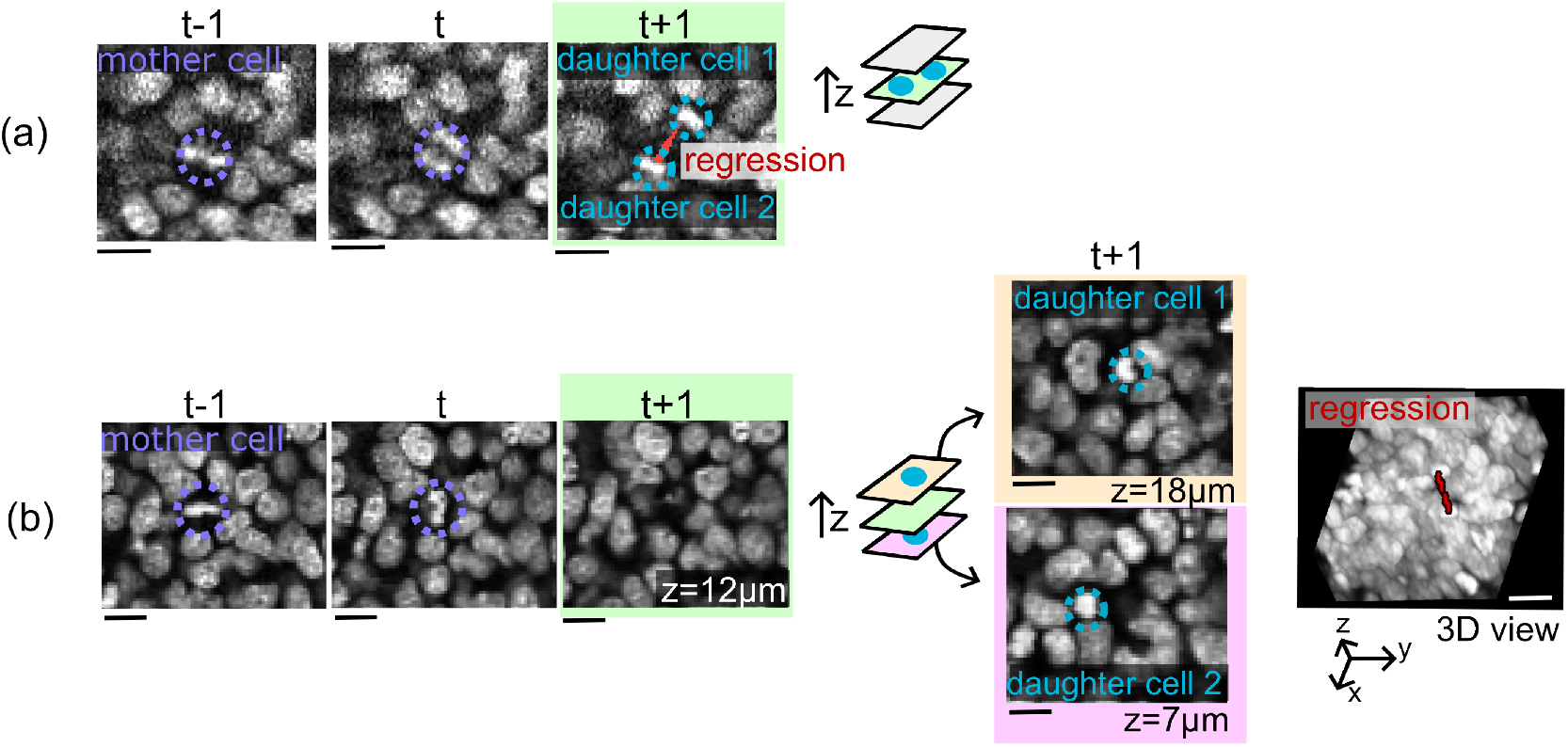
DARE3d nuclei model applied to the gastruloid bulk (Dataset 3): (a) Example of a division in the imaging *xy* plane versus one perpendicular to it, with both daughter cells in the same plane as the mother cell. (b) Example of a division perpendicular to the *xy* plane, with the daughter cells located in the planes above and below, which challenges both annotation and detection. Red lines in (a,b) are the output of the regression code.

**FIG. 5.**
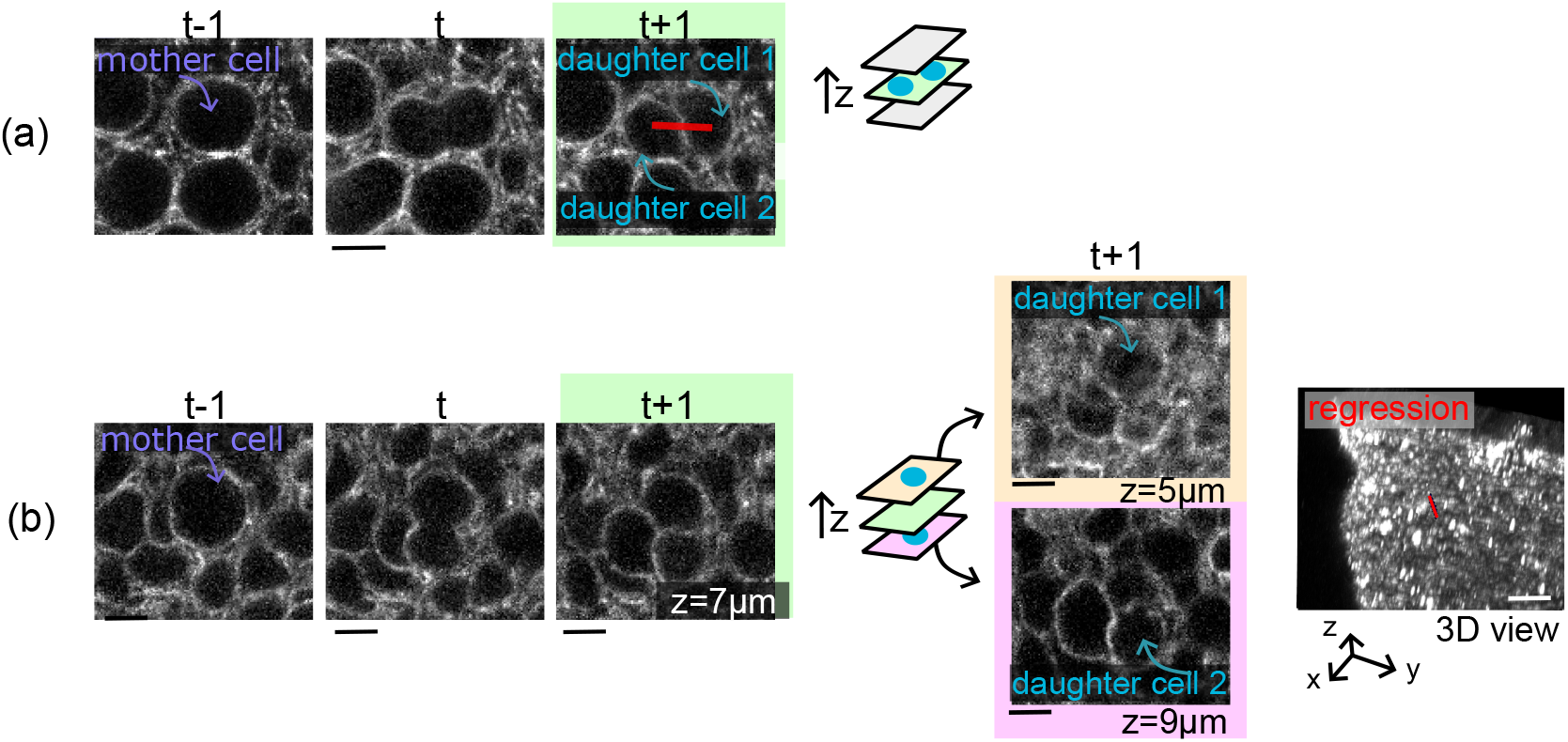
DARE3d membrane model applied to the avian neuroepithelium (Dataset 2): (a) Example of a division in the imaging *xy* plane versus perpendicular to it, with both daughter cells in the same plane as the mother cell. (b) Example of a division perpendicular to the *xy* plane, with the daughter cells located in the planes above and below, which challenges both annotation and detection. Red lines in (a,b) are the output of the regression code.

**FIG. 6.**
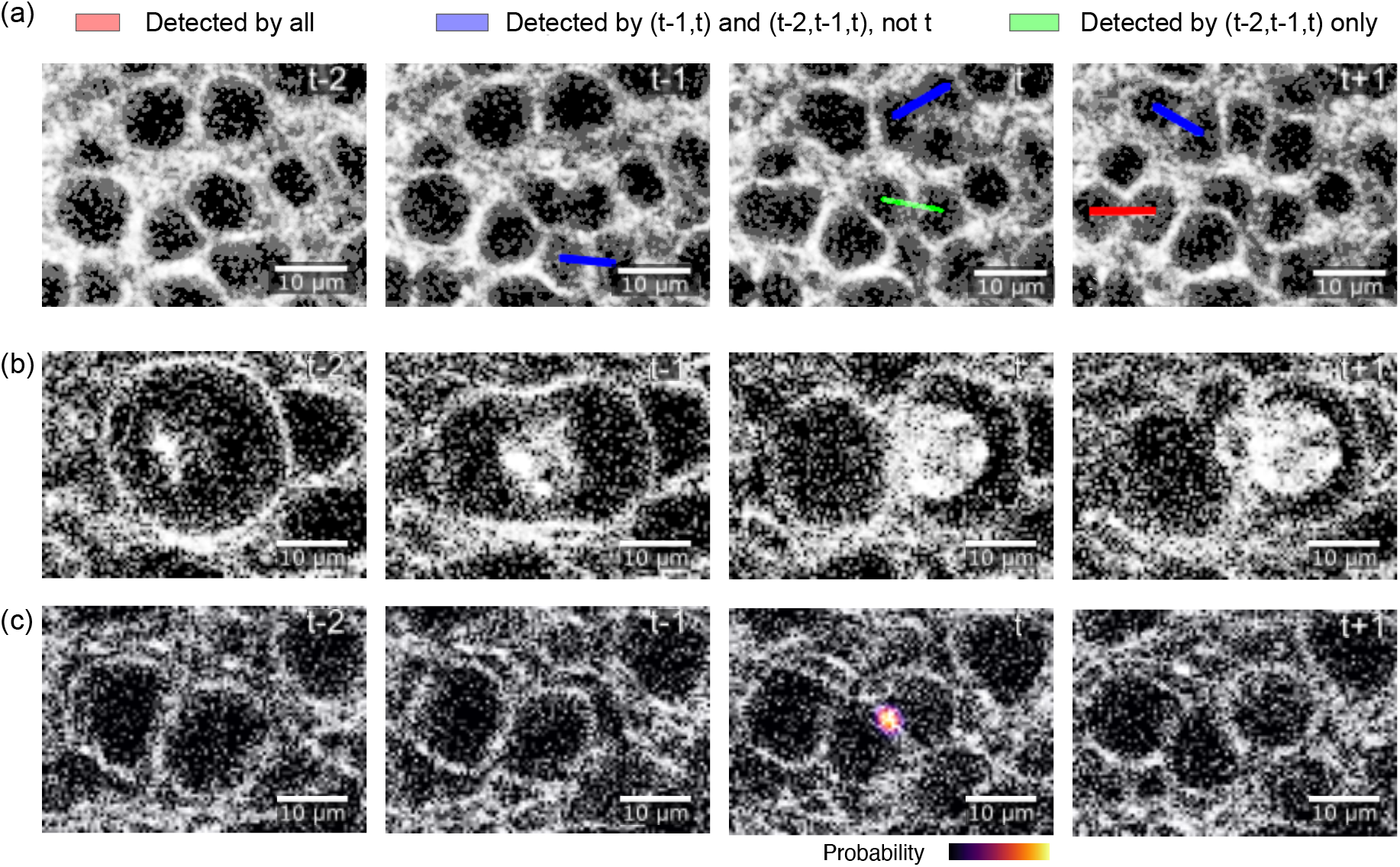
DARE2d membrane model discussion: (a) Example of detected and missed divisions when using a single (*t*), two (*t* − 1, *t*), or three (*t* − 2, *t* − 1, *t*) frames. Red: divisions detected in all three cases. Blue: divisions detected using two (*t* − 1, *t*) and three (*t* − 2, *t* − 1, *t*) frames but missed using *t* only. Green: divisions detected only using three (*t* − 2, *t* − 1, *t*) frames. (b-c) Examples of (b) a missed division and (c) a false detection.

**FIG. 7.**
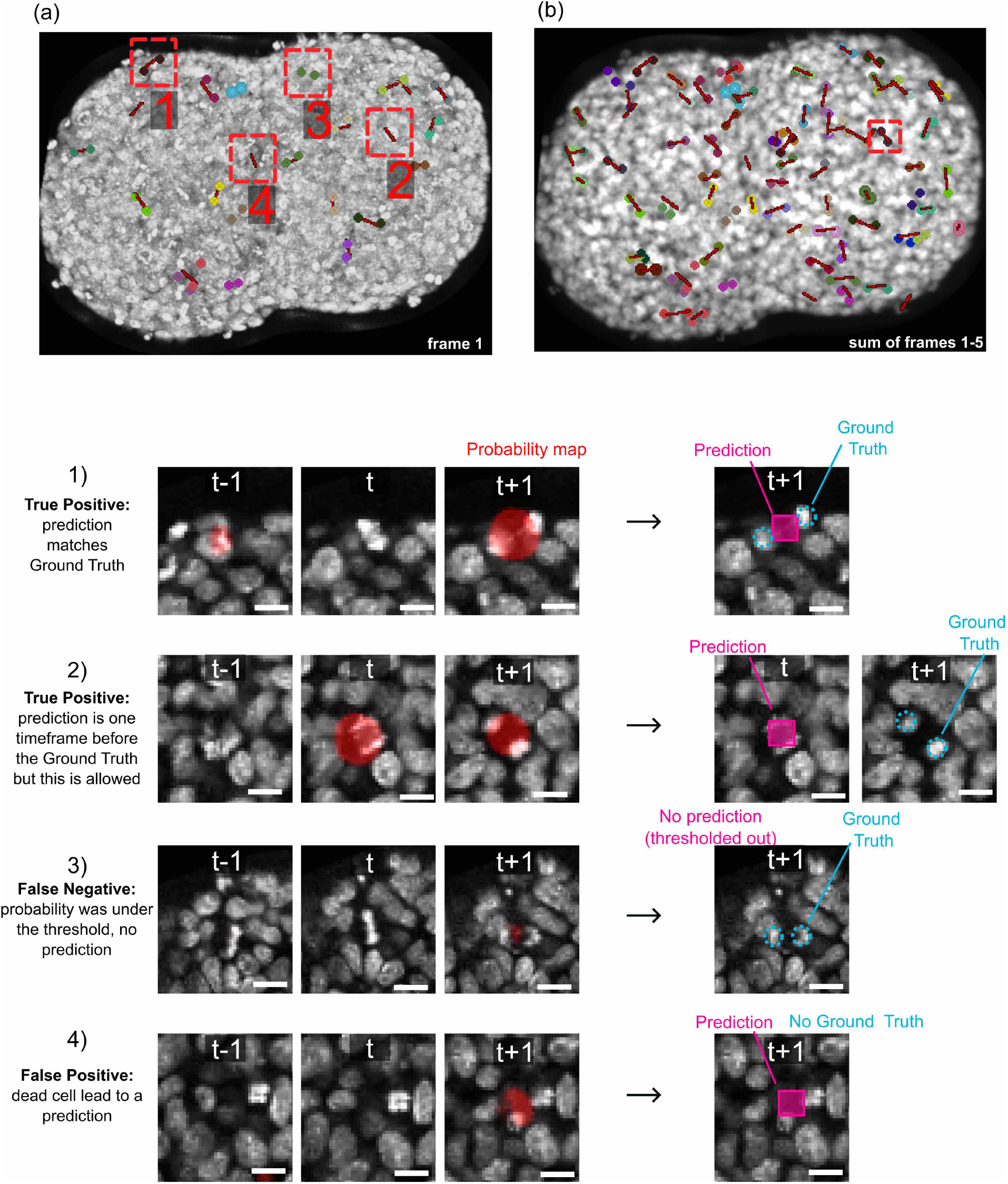
DARE3d nuclei model results on the validation set. (a) 3D projection of one frame with the regression overlaid on the ground truth. Colored dots are daughter cells following division. (1) Example of a True Positive (TP), where the probability map is thresholded into a prediction matching the ground truth. (2) At this frame, the prediction matches no ground-truth annotation because the daughter cells were annotated at the next frame (*t* + 1). A one-frame shift is not counted as erroneous, thanks to our post-processing algorithm defined in Sec. III B 3, resulting in this division being correctly labeled as a TP. In map (b), which is the sum of the subsequent frames, the regression matches the ground truth. (3) Example of a False Negative (FN), where the U-Net wrongly predicts a low probability for these pixels to correspond to a division event. The probability is below the threshold and counts as an absence of prediction. (4) Example of a False Positive (FP): the midpoint segmentation, stage-1 DARE3d nuclei model wrongly predicts a high probability for these pixels to correspond to a division event, whereas it is actually a bright dead cell. Scale bar: 10*µ*m.

A key factor for score optimisation was the standardisation of imaging conditions: spatial and temporal resolution need to remain consistent, so that the network is presented with the same typical sequence of events (e.g. one frame for each of the 3 division phases—prophase, metaphase, anaphase; see Fig. 2) and similar object texture after image renormalisation and isotropisation. In particular, differences in image intensity due to signal loss along the sample were compensated using tools from Ref. [19], to ensure the absence of detection bias along the *z*-stack.

##### Orientation error

A typical one-pixel error on the endpoints of a 30-pixel-long segment in 3D leads to an orientation error of approximately 2^°^. However, annotations are performed on the raw microscopy images and are typically spread over 3–4 frames along the *z* direction, which leads to *±*10-voxel errors in the isotropised movies, mainly along *z*. This implies that a typical mismatch between our predictions and the ground truth is on the order of ≈20^°^; our mean angular error of DARE3d is not significantly larger than 20^°^.

##### Application to a new dataset

We then applied the DARE3d nuclei model to a different experimental condition, namely gastruloids spreading on a substrate (96h old), for which we do not have a ground-truth dataset. Visually, the results appear to agree with expectations (see **Movie** 7). A substantial number of FPs appeared at the propagating front, where the signal-to-noise ratio is extremely low.

#### 2. Discussion: 3D membrane model

##### Localisation errors

Our DARE3d membrane model successfully detects cell divisions occurring in the basal direction (corresponding to lower *z* planes) and captures out-of-plane divisions in neuroepithelium movies. A few FNs arose from cells located at the peripheral edges of the imaging volume, where truncated signals prevent the model from reaching the detection threshold (see **Movie** 9). Regarding FPs, some cases correspond to double-counting events, where a single cell is erroneously localised twice; the post-processing step described in Sec. III B 3 is intended to remove such cases. Further refinement of the post-processing algorithm could potentially suppress these FPs.

##### Orientation error

The discussion is similar to that presented in Sec. V B 1.

### C. Discussion: 2D vs 3D

#### Performance on the segmentation

To compare DARE2d and DARE3d on matched inputs, we focus on the same neuroepithelium recordings in two corresponding formats: a 2D time series (dataset 1), obtained as the average-intensity projection of the associated 3D *z*-stack movie (dataset 2). **Movie** 10 presents a side-by-side comparison of the segmentations produced by DARE2d and DARE3d (Set 1 sequences in dataset 2 corresponding to a portion of Set 5 from dataset 1). Overall, we observe very similar behaviour. Representative cases where either DARE3d or DARE2d fails while the other succeeds are highlighted in **Movie** 10. In one rare instance, DARE2d yields a FP whereas DARE3d does not; since DARE2d benefits from a larger training set, a plausible explanation is that DARE3d rejects this detection by leveraging 3D contextual information.

##### Performance on the regression

The regression angular error is larger in 3D than in 2D; this is mostly due to the lower sampling in the *z* direction than along the imaging plane (see Discussion V B 1). Yet, even on isotropized images, it is inherently more difficult to estimate the orientation of a nematic object in 3D than in 2D; to illustrate this, we define the root-mean-square error *ϕ*_rms_ for a random-guessing detector using the probability density *p*(*ϕ*) of the angular distribution:

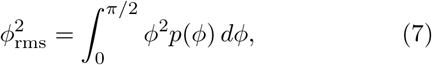

with *p*(*ϕ*) = 2*/π* in 2D, resulting in 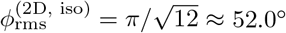, and *p*(*ϕ*) = sin *ϕ* in 3D, which biases the error towards larger angles, giving 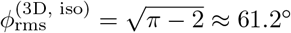. Our 2D and 3D angular errors are well below these values.

## VI. PERSPECTIVES AND CONCLUSION

### Alternative architectures

Our framework relies on human annotations. Self-supervised or unsupervised methods could reduce the need for such manual effort [32, 33]. However, substantial manual intervention during post-processing would likely still be required to ensure accuracy, for instance to distinguish cell divisions from other cellular events such as extrusion.

Our framework is also lightweight, enabling rapid retraining on standard personal computers. Although Transformers [34] could further improve division localisation, current Transformer-based approaches typically still require large amounts of annotated data, which may limit their suitability for rapid retraining on new datasets.

## Conclusion

In this article, we optimized two neural networks for characterizing cell divisions from image sequences, in both 2D and 3D. We also investigated (i) the optimal representation parameters for the cell-division detector and (ii) the influence of preceding image frames around the event on predictive performance.

## AUTHOR CONTRIBUTIONS

S. T. and J.-F. R. initiated and conducted the research. Experimental data were produced by M. S. for the avian neuroepithelium and by S.T. for the mouse gastruloids. M. K., S. T., J.-F. R. and R. K. developed and optimized the neural networks and all authors contributed to data formatting, augmentation, visualisation strategies, and optimization of the training strategies. M. K., M. S., and R. K. contributed to the 2D data annotations; A. G., J.-F. R., S. T., J. V., and Q. S. R. contributed to the 3D data annotations. All authors contributed to the manuscript write-up.

## ETHICS AND COMPETING INTERESTS

We have no competing interests. No ethical approval was required for our work.

## ACKNOWLEDGEMENTS

J.-F. R., M. K. and S. T. acknowledge O. Cochet-Escartin for useful discussions on the angle inference.

The project leading to this publication has received funding from France 2030, the French Government program managed by the French National Research Agency (ANR-16-CONV-0001) and from Excellence Initiative of Aix-Marseille University - A*MIDEX, and by a generic grant to S.T. ANR-22-CE30-0021. This work was supported by the Fondation pour la Recherche Médicale to A. G, grant number FDT202404018538. This work was granted access to the HPC resources of IDRIS under the allocation AD010314339 made by GENCI.

## DATA AVAILABILITY

Here we provide links to

- Zenodo: DARE2d image sequences, annotations and models [35],
- GitHub: DARE2d code [36],
- Zenodo: DARE3d image sequences, annotations and models [37],
- GitHub: DARE3d code [38].

